# Heritable Immunization Establishes a New Model for Pathogen Control

**DOI:** 10.1101/2024.12.19.629026

**Authors:** Joanna Buchthal, Emma J. Chory, Zachary Hill, Yu Zhou, Devanand Bondage, Summer DeAmelio, Julien Freeman, Rudolf Jaenisch, Styliani Markoulaki, Wayne A. Marasco, Sam R. Telford, Kevin M. Esvelt

## Abstract

Heritable immunization represents a promising approach for controlling infectious diseases by embedding immunity directly into the genomes of wild species that spread human pathogens. Here, we report the genetic engineering of *Mus musculus* to produce a neutralizing, protective single-chain antibody against *Borrelia burgdorferi*, the causative agent of Lyme disease. Engineered mice stably produced a LA-2 scFv-albumin fusion protein across multiple generations, demonstrating robust heritability and stability of gene expression. Following sequential challenges with infected and uninfected ticks, heterozygous mice exhibited strong resistance to infection, effectively interrupting the *Borrelia burgdorferi* disease transmission cycle. Having recently established novel protocols to genetically engineer the white-footed mouse, *Peromyscus leucopus*, a key reservoir of Lyme disease, these findings demonstrate the feasibility of heritable immunization as a potential strategy for mitigating Lyme disease transmission in the environment. More broadly, engineered reservoir immunity may offer a generalizable approach to controlling vector-borne and zoonotic disease with profound potential to improve human health.

## INTRODUCTION

Heritable immunization—the encoding of pathogen resistance into the genome of organisms—offers a novel strategy for controlling infectious diseases by integrating immune protection directly into the germline of species that transmit pathogens. This strategy enables the stable transmission of immunity across multiple generations by providing continuous, systemic protection against pathogens in reservoir hosts. One potential approach, encoding neutralizing antibodies within the genome directly, has the potential to disrupt disease transmission cycles at their source, particularly in well-characterized host–pathogen systems. Unlike conventional vaccination, which requires repeated administration to each generation, engineering reservoir species could synergize with concomitant interventions by providing a perpetuating mode of reducing enzootic transmission of infections, particularly those maintained by rodents.

Lyme disease, the most common vector-borne disease in the United States, is a multisystem infection with a global public health impact. The transmission cycle involves a complex interplay between tick vectors, reservoir hosts, and the environment (Radolf et al., 2012). Infection is not vertically transmitted; rather, infections are transmitted between each new generation of reservoirs and tick vectors. On the East Coast of the United States, *Peromyscus* species, particularly the white-footed mouse (*Peromyscus leucopus*), serve as the primary reservoir for *Borrelia burgdorferi* (Bunikis et al., 2004). In Europe, the bank vole (*Myodes glareolus*) and the wood mouse (*Apodemus sylvaticus*) are key reservoirs for *Borrelia afzelii* (Lindsø et al., 2024), while in Asia, the striped field mouse (*Apodemus agrarius*) serves as a reservoir of *Borrelia garinii* (Zhang et al., 2010). Additionally, *Mus* species, including *Mus musculus*, have been identified as potential reservoirs of *Borrelia* in Europe (Frandsen et al., 1995).

Given the key role played by rodents in the transmission of Lyme disease, many prevention strategies target these hosts to disrupt the transmission cycle. For example, “tick control tubes” have been deployed to reduce tick populations and interrupt transmission by distributing permethrin-treated cotton, which mice incorporate into their nests (Deblinger & Rimmer, 1991; Stafford, 1991). Several additional strategies targeting rodent reservoirs focus on Borrelia’s major outer surface protein A (OspA), a key protective antigen expressed by *B. burgdorferi sensu lato* (s.l.). Immunization with OspA induces an antibody response that provides protection by a unique mode of action: anti-OspA antibody ingested by an infecting tick is thought to incapacitate or destroy bacterial spirochetes in the tick prior to their attaining infectivity, thereby preventing transmission (Fikrig et al., 1992). Parenteral immunization of wild white-footed mice with an OspA subunit vaccine in field studies demonstrated a reduction in the prevalence of infected ticks (Tsao et al., 2004). Oral vaccination with recombinant OspA induces a protective response in mouse models (Fikrig et al., 1991), and a commercially available baited rOspA vaccine has been field tested and may be effective in reducing *B. burgdorferi* transmission in wild mice (Richer et al., 2014).

Building on the success of previous OspA-based immunization strategies, we explored the use of genetic engineering to alter the reservoir capacity of key hosts by encoding OspA-targeting antibodies in the mouse genome. This approach, inspired by vector-borne disease control strategies such as the genetic engineering of mosquitoes to combat malaria (Hammond et al., 2016; Hoermann et al., 2022), introduces the novel concept of heritable immunization—embedding antibody-based immunity into the germline to achieve lasting resistance. While mice have been engineered to produce human antibodies, such as the XenoMouse® (Abgenix, Inc., Fremont, CA) and HuMAb Mouse® (GenPharm-Medarex, San Jose, CA), these models are used for therapeutic monoclonal antibody discovery and production. They do not produce preexisting, pathogen-specific antibodies, nor are they designed to confer lifelong resistance to specific infections or transmit immunity to offspring (Jakobovits et al., 2007; Peterson, 2005). Unlike prior research where engineered mouse mothers transferred disease resistance to their pups through antibodies in breast milk (Castilla et al., 1998), our approach aims to establish continuous systemic protection in the reservoir species, persisting throughout their lifespan, and lasting for many generations in the environment.

In this study, transgenic *Mus musculus* were engineered to express LA-2, a well-characterized protective monoclonal antibody derived from *Mus* that targets *Borrelia burgdorferi* (Schaible et al., 1990), with the aim of disrupting Lyme disease transmission. Initial *in vitro* experiments were conducted to optimize genetic engineering efficiency, antibody design and stabilization via leader sequences, and bicistronic elements. Our original goal was to achieve liver-specific expression, but the initial full-length LA-2-expressing models exhibited insufficient immunity. This led to the development of a mouse model in which the LA-2 antibody was reformatted into a single-chain variable fragment (scFv), fused to albumin for enhanced stability and constitutive, ubiquitous expression from the Rosa26 locus. These engineered mice exhibited robust antibody production across multiple generations, and demonstrated strong immune protection, leading to a statistically significant reduction in the transmission of disease-causing bacteria following exposure to infected ticks. These results establish heritable immunization as a viable strategy for the prevention of diseases with mammalian reservoir species.

## RESULTS

### Testing Albumin Expression Machinery

To develop a heritable immunization strategy against Lyme disease, we began by optimizing the expression of the anti-*Borrelia burgdorferi* antibody LA-2 in mouse hepatocytes. To achieve tissue-specific antibody release into the bloodstream, we co-opted the expression machinery of the native albumin gene and evaluated its ability to drive protein expression. Specifically, we investigated whether the minimal albumin promoter was sufficient to support protein expression when inserted proximal and divergent to the endogenous bi-directional albumin enhancer or if an additional albumin enhancer was necessary (Pinkert et al., 1987).

We generated two distinct CRISPR-modified Hepa 1-6 cell lines: 1) the first expressing tdTomato from the minimal promoter alone and 2) a second which incorporated the albumin enhancer (**Supp Fig. 1A**). Briefly, integration cassettes were co-transfected into Hepa 1-6 cells along with SpCas9 and guide RNA targeting sequences ∼300 bp proximal to the albumin enhancer. Following hygromycin selection, integration was confirmed by PCR. RT-qPCR analysis revealed comparable levels of tdTomato and native albumin mRNA expression in the absence of the additional enhancer (**Supp Fig. 1B**), suggesting that the minimal promoter alone would be sufficient for robust mRNA expression.

### Leader Sequence Selection

We next aimed to identify a leader sequence that could efficiently direct the anti-*Borrelia burgdorferi* LA-2 antibody for secretion into the bloodstream. To optimize LA-2 secretion, we evaluated potential leader sequences using both *in silico* and *in vitro* approaches. Using IMGT’s V-Quest tool, we identified candidate leader sequences based on their homology to the LA-2 heavy chain variable domain, and additionally analyzed leader sequences from albumin, alpha-fetoprotein, and fibronectin (**Supp Fig. 1C**). Cleavage efficiency was assessed using SignalP software to predict the likelihood of proper cleavage between the leader and LA-2 antibody sequence (**Supp Fig. 1C**). From these analyses, two leader sequences were selected for further testing. To assess antibody production from the selected leaders, we designed three expression vectors containing 1) a reformatted version of LA-2 as an scFv-Fc fusion protein and 2) either one of the two selected leader sequences, or 3) no leader sequence (**Supp Fig. 1D**). These constructs were transfected into Hepa 1-6 cells, and LA-2 scFv-Fc secretion was quantified by ELISA over 48 hours, using a secondary antibody specific to IgG2b for detection. Both leader sequences resulted in expression, with one yielding slightly higher levels. This sequence was therefore selected for all transgenic mouse designs (chosen leader: MAWVWTLLFLMAAAQIQA) (**Supp Fig. 1E**).

### Bicistronic Element Selection

To achieve expression of the full-length LA-2 antibody, we utilized bicistronic elements commonly employed in antibody expression cassettes to produce both heavy and light chains without additional expression machinery. We evaluated several 2A peptides and internal ribosome entry sites (IRES) to determine which element best facilitated the coordinated expression of both chains. Each construct featured a CMV promoter driving expression of the heavy chain, then a distinct 2A sequence or an IRES, and finally the light chain. These constructs were transfected into Hepa 1-6 cells and supernatant was collected 72 hours post-transfection for analysis. IgG2a ELISAs were performed to quantify antibody concentration, while OspA ELISAs assessed binding to the *Borrelia* surface protein. T2A was chosen from the elements tested (**Supp Fig. 1F**).

### Design Validation and Guide Testing

To finalize our design and identify the optimal guide for targeting the albumin locus, we generated four CRISPR-modified cell lines incorporating key design elements from previous constructs along with the full-length LA-2 antibody. To quickly obtain near-clonal isolates, a selectable marker was introduced to facilitate the isolation of LA-2-positive cell populations (**Supp Fig. 1G**). Hepa 1-6 cells were co-transfected with the construct encoding full-length LA-2, SpCas9, and one of four guide RNAs targeting the region 300 bp proximal to the endogenous albumin enhancer. After three weeks of puromycin selection, we established four populations of LA-2-positive, puromycin-resistant Hepa 1-6 cells. We performed RT-qPCR to measure LA-2 mRNA levels, along with albumin and the selectable marker, using five distinct qPCR primer pairs, with expression normalized to the housekeeping gene β-actin (**Supp Fig. 1H**). All lines expressed LA-2 mRNA, with guide 3 yielding substantially higher levels, presumably due to more efficient homozygous insertion. We then evaluated antibody production by performing an IgG2a ELISA on the supernatant from the four LA-2 CRISPR cell lines. One line (from guide 3) measurably secreted IgG2a (**Supp Fig. 1I**).

### Generation and Validation of Liver-Specific Full-Length LA-2 Expressing Mice

To generate mice expressing the full-length LA-2 antibody, we next constructed a vector containing the aforementioned components, but without a selection marker (**Fig. 1A**). We performed pronuclear injection to introduce SpCas9, guide 3, and the full length LA-2 expression cassette into BDF1 mouse embryos. Of the 45 pups delivered, sequencing confirmed that two carried a complete LA-2 antibody knock-in, with one founder exhibiting the correct insertion at the target locus.

**Figure 1.**
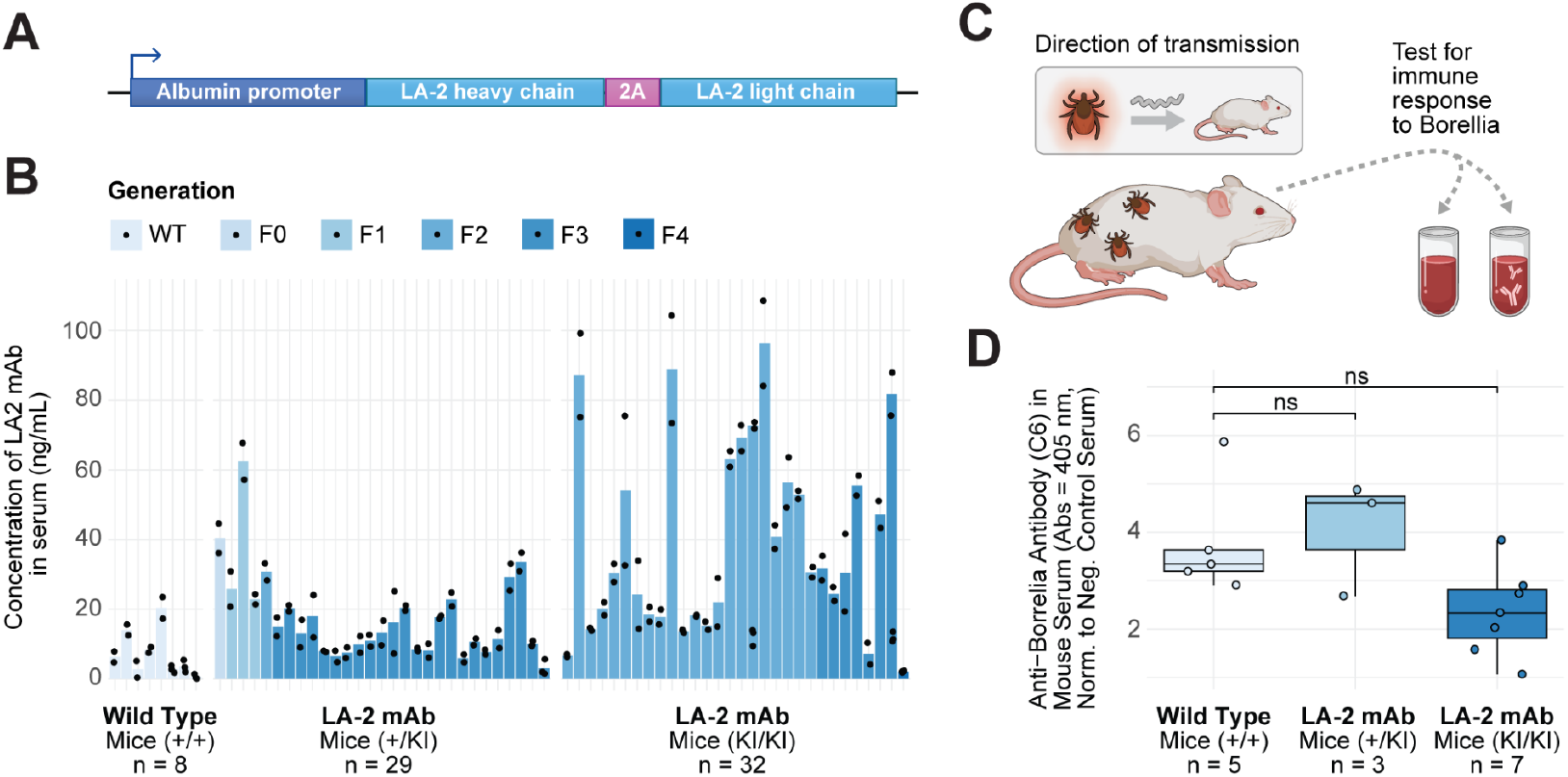
Generation, Validation, and Infection Challenge of Liver-Specific Full-Length LA-2-Expressing Mice. **A)** Schematic of the construct introduced into the mouse genome, 300 bp proximal to the native albumin enhancer. The minimal albumin promoter drives the expression of the full-length LA-2 monoclonal antibody. **B)** Quantification of LA-2 antibody concentration in the serum of heterozygous, homozygous, and wild-type mice across multiple generations (F0 through F4). **C)** Diagram of the tick challenge. Uninfected engineered and control mice were exposed to *Borrelia burgdorferi*-infected ticks, and post-infestation serum was evaluated for anti-*Borrelia* antibodies to determine infection status. **D)** Endpoint ELISA results from transgenic and wild-type mice 21 days post-infestation, detecting *B. burgdorferi*-specific antibodies against C6, a sensitive marker of infection.

Engineered mice were bred for multiple generations to assess both the heritability and stability of LA-2 antibody expression in homozygous and heterozygous mice. OspA ELISAs were then performed to quantify the concentration of LA-2 antibodies that correctly bound the protective epitope on OspA (**Fig. 1B**). However, antibody levels showed major differences, with homozygous mice exhibiting the highest variability, ranging from negligible wild-type-equivalent levels to over 80 ng/mL.

### Testing the Susceptibility of Liver-Specific Full-Length LA-2 Expressing Mice to Infection

To evaluate the efficacy of liver-specific, full-length LA-2 antibodies in conferring heritable resistance to *Borrelia burgdorferi* infection, we challenged engineered mice from different filial generations and genotypes with *Borrelia*-infected *Ixodes dammini* nymphs (**Fig. 1C**). Infected nymphs were prepared by feeding larval *Ixodes dammini* on *Mus musculus* infected with the low-passage N40 *Borrelia burgdorferi* strain (wildtype), which we maintain through tick-mouse-tick transfer (Fikrig et al., 1992). All infected nymphs were derived from a single vial with a 70% infection rate, as determined by indirect immunofluorescence (data not shown). During infestation, 10 nymphs were applied to the ears and nape of the neck of each anesthetized mouse. Nymphs were collected 4-6 days post-infestation. The number of engorged ticks recovered varied from 2 to 7 per mouse. Mice were held for 21 days post-infestation, after which serum was collected and analyzed using an endpoint ELISA to detect *B. burgdorferi*-specific antibodies against C6, a sensitive marker of infection (Bacon et al., 2003). Although some homozygous mice exhibited very low or undetectable C6 antibody levels, suggesting potential immunity, neither the homozygous nor heterozygous mice showed a statistically significant reduction in anti-C6 antibody levels compared to wild-type controls. As a result, this line was discontinued due to insufficient immunity (**Fig. 1D**).

### Generation of Rosa26-Targeted LA-2 scFv-Albumin Mice

To develop a mouse model with enhanced antibody expression and improved resistance to infection, we next generated a mouse with key modifications to the original design. First, we reformatted the full-length LA-2 antibody into a single-chain variable fragment (scFv) to address the expression challenges associated with bicistronic elements and to prevent the mispairing of the full length LA-2 heavy and light chains with endogenous antibodies. During reformatting, we tested binding and observed an almost one-log reduction in affinity when converting the full-length LA-2 IgG to an scFv with a Vh-(G4S)3-Vl linker (**Supp Fig. 2**).

To optimize LA-2 scFv expression, stability, and binding, we evaluated 15 different linker sequences by generating expression constructs with a unique linker positioned between the LA-2 scFv-Fc heavy and light chains (**Supp Fig. 3A**). Three constructs employed multimers of the commonly used GGGGS pentapeptide, while the remaining linkers were designed to modulate flexibility and length by varying the number of glycine and serine residues. To assess the various linkers, constructs were transfected into Lenti-X 293T cells, and 72 hours post-transfection, supernatant was collected for OspA and IgG2a ELISAs. The optimal linkers were identified by comparing binding data to that of purified full-length LA-2 antibody, with the top-performing linker demonstrating superior binding characteristics. This linker sequence, GGGGSGGGGSGGGGSGGGG, was selected for mouse model generation (**Supp Fig. 3B**).

To generate engineered mice, we next developed a targeting construct with modifications aimed at enhancing antibody expression and stability. Given the inherently short half-life of scFvs, we fused the LA-2 scFv to albumin, a strategy previously explored in therapeutic contexts to improve protein stability and extend half-life (Marsh & Owen, 2023; Tao et al., 2021). To ensure robust antibody production, we utilized the CAG enhancer/promoter, known for driving high gene activity across various loci, including Rosa26 (Gu et al., 2018) (Gu et al., 2018). We integrated the LA-2 scFv-albumin into the *Mus* Rosa26 locus, a well-established safe harbor site known for stable and consistent transgene expression (**Fig. 2A**).

**Figure 2.**
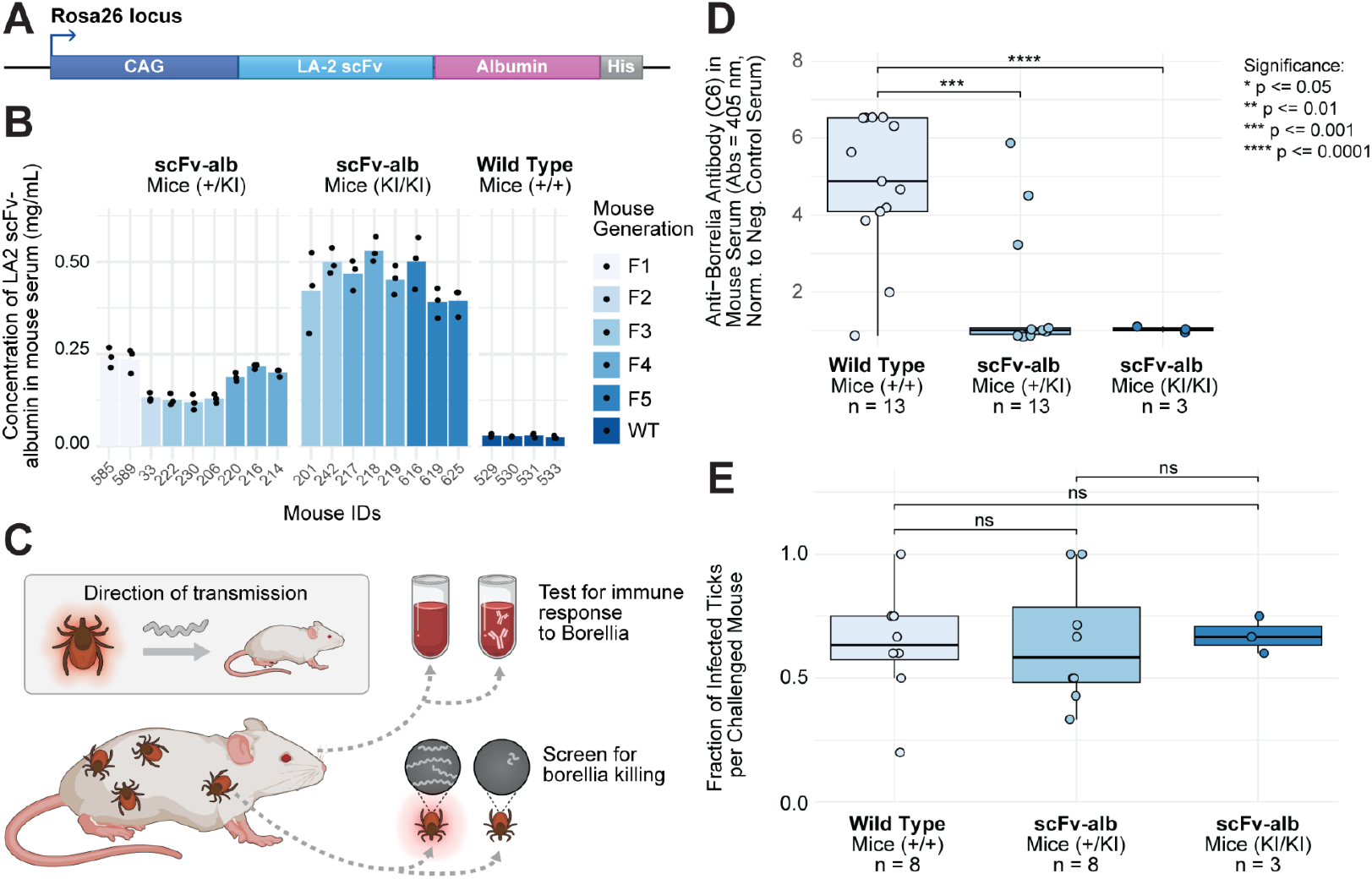
Generation, Validation, and Infection Challenge of LA-2 scFv-Albumin Expressing Mice. **A)** Schematic representation of the transgene inserted into the Rosa26 locus, encoding the LA-2 scFv fused to mouse albumin under the control of the CAG promoter. **B)** Serum concentrations of LA-2 scFv-albumin in heterozygous and homozygous mice across multiple generations, measured by ELISA. **C)** Illustration of the tick challenge: Uninfected engineered and control mice were exposed to *Borrelia burgdorferi*-infected ticks. Post-infestation, serum was analyzed for signs of infection, and ticks were evaluated for infection status. **D)** Endpoint ELISA results from transgenic and wild-type mice 21 days post-infestation, detecting *B. burgdorferi*-specific antibodies against C6, a sensitive marker of infection. **E)** Graph showing the fraction of *Borrelia*-infected ticks per individual mouse, comparing ticks that fed on transgenic versus wild-type mice. Engorged ticks were collected, stored, and assessed for the presence of *Borrelia* spirochetes using indirect immunofluorescence.

We performed pronuclear injection to introduce SpCas9, guide RNA, and the LA-2 scFv-albumin expression cassette into embryos from B6 mice. Eighteen pups were produced and tested, with only one showing a complete antibody knock-in at the correct locus.

### Testing Heritable Antibody Expression in Rosa26-Targeted LA-2 scFv-Albumin Mice

Engineered mice were bred for six generations, with each generation genotyped to ensure the presence of the LA-2 scFv-albumin antibody gene. Antibody expression was evaluated by testing serum samples from each generation for OspA-binding LA-2 antibody using a direct ELISA with an anti-albumin secondary antibody. Engineered mice produced at least 1000-fold more antibody compared to the previous design, with homozygous mice expressing roughly twice the amount seen in heterozygotes (**Fig. 2B**). Unlike the previous design, there was no variability in antibody production; animals with one copy of the gene consistently produced close to 0.25 mg/mL of antibody, while those with two copies produced approximately 0.5 mg/mL. Antibody expression remained consistent across successive generations, confirming stable, heritable expression.

### Testing the Susceptibility of Rosa26-Targeted LA-2 scFv-Albumin Mice to Infection

To evaluate the resistance of newly engineered LA-2 scFv-albumin-expressing mice to *Borrelia burgdorferi* infection, we challenged these mice alongside wild-type controls with *B. burgdorferi*-infected nymphs (**Fig. 2C**). As in the previous challenge, ten nymphs were applied to each mouse and allowed to feed to repletion. Twenty-one days post-challenge, mice were bled, and serum was analyzed for *B. burgdorferi*-specific IgG antibodies against the C6 peptide (**Fig. 2D**). Both homozygous and heterozygous mice showed a statistically significant reduction in anti-C6 antibody production compared to wild-type controls. Homozygous mice, in particular, demonstrated robust protection from infection. Heterozygous mice also exhibited significant immunity (p=1.8E-4). The small subset showing reduced protection may reflect differences in infected tick or *Borrelia* burdens. While further studies are required to fully characterize the level of protection in both heterozygous and homozygous mice, these results indicate a strong and heritable immune response that is protective against tick challenge, especially in homozygotes.

### Evaluating Tick Infection Following the First Challenge with Rosa26-Targeted LA-2 scFv-Albumin Mice

Next, the nymphs from the previous challenge were assessed for infection to evaluate the borreliacidal activity of the scFv-albumin antibodies in the engineered mice, following a previously described protocol (Fikrig et al., 1992). To assess infection, engorged nymphs were collected 4-6 days post-infestation and stored for 14 days before analysis (Fikrig et al., 1992). Ticks were homogenized, applied to slides, and stained by indirect immunofluorescence using a rabbit polyclonal serum against *B. burgdorferi*. Slides were examined at 400x magnification under epifluorescence, and the stained homogenates were scored for the relative density of *B. burgdorferi* and spirochetal morphology (intact vs. disrupted). For analysis, each tick was categorized based on the presence or absence of spirochetes, regardless of infection intensity (**Fig. 2E**). Intriguingly, while the scFv-albumin antibodies provided protection to the mice, they did not appear to reduce the spirochete load in the infected ticks used to challenge those mice, contrary to the previously reported mode of action for anti-OspA antibodies (Fikrig et al., 1992).

### Testing *Borrelia* Transmission to Uninfected Ticks Feeding on Previously Challenged Rosa26-Targeted LA-2 scFv-Albumin Mice

Even without *Borrelia* clearance, engineered mice may reduce or block transmission of the infection to the next generation of larvae. To assess the infection status of the mice post-challenge and determine whether anti-OspA antibodies impacted *Borrelia* transmission, we performed xenodiagnosis using uninfected larval ticks (**Fig. 3A**) (Marques et al., 2014). Twenty-one days after the challenge with infected ticks, mice were infested with uninfected larval *Ixodes dammini* ticks. Engorged larvae were collected and maintained until molting, approximately 4-5 weeks later. Post-molt, ticks were dissected and their gut contents smeared onto slides and stained via indirect immunofluorescence using a rabbit polyclonal serum specific to *Borrelia burgdorferi*. A minimum of 50 fields (X320) were examined per slide before a negative result was declared, and the infection rates per mouse were quantified (**Fig. 3B**). Detection of spirochetes in any tick (e.g., 1 out of 5) was considered evidence of an infected and infectious mouse. A comparison of ticks feeding on wild-type versus engineered mice revealed a highly significant difference in infection status (p=6.9E-4). Moreover, results indicate that 80% of the previously challenged engineered mice were completely free of infection.

**Figure 3.**
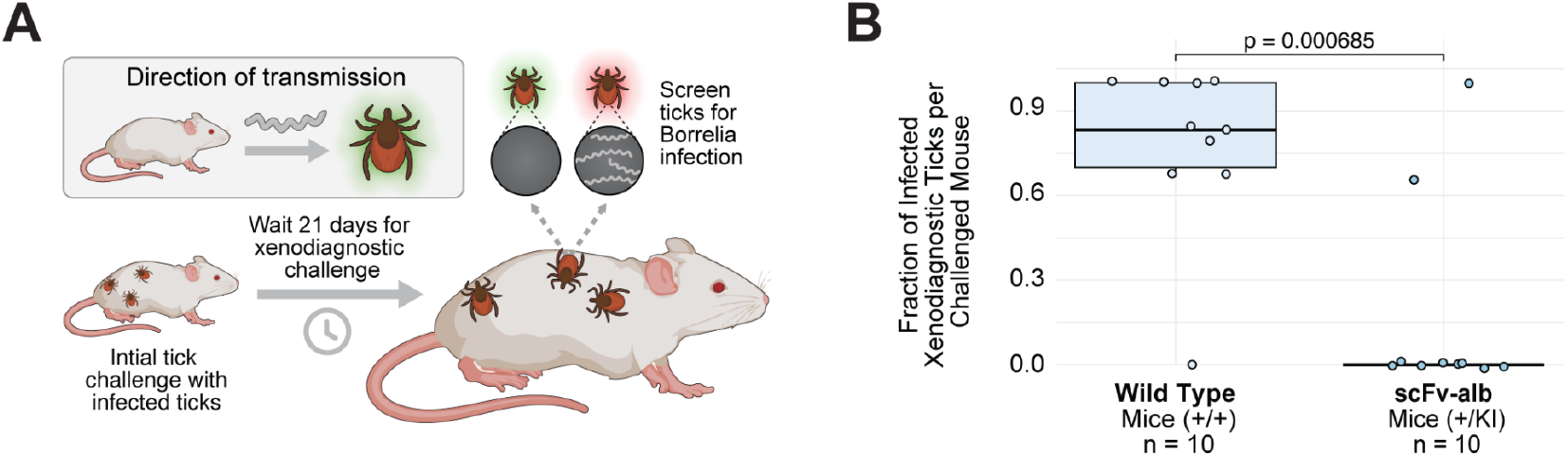
Xenodiagnostic Challenge and Infection Analysis in LA-2 scFv-Albumin Expressing Mice. **A)** Schematic of the xenodiagnostic challenge. Uninfected larval ticks were fed on previously challenged Rosa26-targeted LA-2 scFv-albumin mice 21 days post-challenge. **B)** Fraction of infected ticks per mouse, assessed by indirect immunofluorescence of dissected tick guts post-molt. Detection of *Borrelia burgdorferi* spirochetes in any tick was considered evidence of an infected and infectious mouse.

## DISCUSSION

By encoding anti-Lyme antibody genes into the germline of *Mus musculus*, we conferred heritable resistance to Lyme disease. Antibody genes passed across multiple generations provided robust immunity to infection. Our results highlight the potential of heritable immunization as a strategy for controlling infectious diseases by genetically engineering reservoir species. These findings have immediate translational relevance, particularly in regions where mouse species are key reservoirs, such as Europe.

The same approach to engineering heritable immunity could be adapted to target a broad spectrum of vector-borne and zoonotic diseases by introducing specific antibodies or immune-modulating factors into the genomes of key wildlife species. Examples include carriers of hantaviruses (Mills et al., 1999), leptospirosis (Gomes-Solecki et al., 2017), Lassa fever (Smither et al., 2023), and other emerging pathogens. Encoding immune protection directly into the germline could advance global efforts to prevent and control infectious diseases impacting animal and human health.

While our results demonstrate effective immune protection, the precise mechanism remains unclear and warrants further study. Previous work suggested that anti-OspA antibodies directly neutralize *Borrelia* in the tick midgut via direct cellular lysis or complement activation (Fikrig et al., 1992). However, the single-chain antibody we employed did not clear *Borrelia* within the tick, suggesting that an alternative mechanism must be responsible for disrupting the transmission cycle. One plausible explanation, as proposed by Wang et al. (2016), is that anti-OspA antibodies interfere with *Borrelia*’s gene switch from OspA to OspC, which is essential for spirochete infection of vertebrates (Wang et al., 2016). This switch, regulated by environmental signals (temperature, pH, and spirochete density) within the tick during a blood meal, is crucial for the successful infection of a mammalian host (de Silva et al., 1999; Schwan et al., 1995). By binding to the surface of *Borrelia*, anti-OspA antibodies may prevent the bacteria from reaching the critical density required for the OspA/OspC gene switch, or alternatively, they may block interactions essential for transmission (de Silva et al., 1999). Without Fc-mediated effector functions, the LA-2 scFv appears sufficient to prevent *Borrelia* from establishing infection in the mammalian host, highlighting that antibody binding alone, independent of Fc-driven immune mechanisms, could effectively disrupt the transmission cycle. Differences in affinity and avidity between the scFv and the full-length antibody may further contribute to these distinct findings. Future iterations incorporating new antibody formats designed to enhance affinity and avidity could be explored for increased efficacy, particularly in applications requiring bactericidal activity or targeting other pathogens.

While our findings represent a critical first step toward heritable immunization, several challenges remain before implementation. Extensive field trials are necessary to assess ecological fitness and efficacy in natural environments, as fitness costs, though not yet observed in the lab, could manifest in the wild. Additionally, the complex ecology of *Borrelia burgdorferi* and the potential role of additional reservoirs (Goethert et al., 2024) suggest that engineering a single species may not fully disrupt transmission in all environments, potentially requiring application of this strategy across multiple hosts. However, our recent advances in rodent ovulation tracking open new avenues for engineering non-model species involved in *Borrelia* transmission, paving the way for a comprehensive strategy for disease control.

This study represents an important advance in genetic immunization, demonstrating the feasibility and potential of heritable immunization strategies in reservoir species. Future research should focus on refining this approach, elucidating the mechanisms of immune protection, and conducting field trials to evaluate the real-world impact of heritable immunization strategies. The integration of such strategies into broader public health initiatives could transform infectious disease control, particularly in regions where vector-borne diseases threaten human and animal health.

## AUTHOR CONTRIBUTIONS

**JB**: Conceptualization, Methodology, Validation, Formal Analysis, Investigation, Writing-Original Draft, Visualization, Project Administration, Funding Acquisition; **EJC**: Methodology, Validation, Writing - Review & Editing, Visualization, Supervision; **ZH**: Investigation; **YZ**: Investigation, Writing - Review & Editing; **DB**: Investigation; **SD**: Investigation; **JF**: Investigation; **RJ**: Writing - Review & Editing, Supervision; **SM**: Supervision; **WAM**: Methodology, Validation, Writing - Review & Editing; **SRT**: Methodology, Validation, Investigation, Resources, Writing - Review & Editing, Supervision; **KME**: Conceptualization, Methodology, Writing - Review & Editing, Supervision, Funding Acquisition.

## ETHICS STATEMENT

The Massachusetts Institute of Technology’s Committee on Animal Care approved all mouse procedures and all mice were maintained at MIT (Cambridge, MA, USA) in strict accordance with all institutional protocols, DOD guidelines and the Guide for the Care and Use of Laboratory Animals. The Tufts University Institutional Animal Care & Use Committee approved all mouse studies performed at Tufts (North Grafton, MA, USA).

## METHODS

### Testing Albumin Expression Machinery

Hepa 1-6 cells were co-transfected with two integration cassettes: one containing the minimal albumin promoter driving tdTomato expression and the other incorporating the rat albumin enhancer (Postic et al., 1999). Transfections were carried out using Lipofectamine 2000 (Thermo Fisher) and Opti-MEM (Gibco), following standard protocols. SpCas9 and one of four CRISPR guides, targeting sequences approximately 300 bp upstream of the albumin enhancer, were included in the transfection mix. The guide sequences used were as follows:

gRNA 1: GAGCTAACCTTCTGTCCTAG

gRNA 2: GCCTTAGCCAGTGTTTGCAC

gRNA 3: GCCTGTGCAAACACTGGCTA

gRNA 4: GCTGGCTAAGGCATGAACTT

Following transfection, cells were cultured under hygromycin selection until only cells with integrated cassettes remained. After 3 weeks of selection, successful integration was confirmed by PCR. RNA was extracted using TRIzol Reagent (Thermo Fisher), and cDNA synthesis was performed using the Quantitect Reverse Transcription Kit (Qiagen). Expression levels of tdTomato and native albumin mRNA were quantified by RT-qPCR using the SensiFAST SYBR Hi-ROX Kit (Bioline), with mRNA levels normalized to β-actin. The following primer sets were used for amplification:

Albumin 1:

Alb-2-qPCR-F: 5’-GACGTGTGTTGCCGATGAGT-3’

Alb-2-qPCR-R: 5’-GTTTTCACGGAGGTTTGGAATG-3’

Albumin 2:

Alb-3-qPCR-F: 5’-TCCAAACCTCCGTGAAAACTATG-3’

Alb-3-qPCR-R: 5’-TGTGTTGCAGGAAACATTCGT-3’

tdTomato 1:

tdTomato-1-qPCR-F: 5’-CTTGTACAGCTCGTCCATGC-3’

tdTomato-1-qPCR-R: 5’-AACTGCCCGGCTACTACTAC-3’

tdTomato 2:

tdTomato-3-qPCR-F: 5’-CGCGCATCTTCACCTTGTAG-3’

tdTomato-3-qPCR-R: 5’-GCGTGATGAACTTCGAGGAC-3’

β-Actin:

Bact-1-qPCR-F: 5’-GGCTGTATTCCCCTCCATCG-3’

Bact-1-qPCR-R: 5’-CCAGTTGGTAACAATGCCATGT-3’

### Leader Sequence Selection

Leader sequences for efficient secretion of the anti-*Borrelia burgdorferi* LA-2 antibody were selected through a combination of in silico analysis and in vitro validation. Candidate sequences were identified using IMGT’s V-Quest tool based on homology to the LA-2 heavy-chain variable domain. Leader sequences from mouse albumin, alpha-fetoprotein, and fibronectin were also evaluated. Cleavage efficiency was predicted using SignalP software.

Three expression vectors were constructed: two containing the selected leader sequences driving expression of the LA-2 single-chain variable fragment fused to the Fc region (scFv-Fc), and one control vector without a leader sequence. The following two leader sequences were tested:

IGHV9-2-1*01: MAWVWTLLFLMAAAQIQA

IGHV9-3*02: MDWLWNLLFLMAAAQIQA

Hepa 1-6 cells were transfected with the aforementioned constructs using Lipofectamine 2000 (Thermo Fisher Scientific) according to the manufacturer’s protocol. Supernatants were collected at 0, 24, and 48 hours post-transfection.

LA-2 scFv-Fc secretion was quantified using a mouse IgG2b ELISA kit (Bethyl Laboratories). ELISAs were performed on the collected supernatants following the manufacturer’s instructions.

### Bicistronic Element Selection

To optimize the co-expression of the heavy and light chains of the full-length LA-2 antibody, five expression constructs were designed. Each construct contained a CMV promoter driving the LA-2 heavy chain, followed by either one of four 2A peptide sequences (F2A, E2A, P2A, or T2A) or an internal ribosome entry site (IRES CVEB) to facilitate co-expression of the light chain.

Hepa 1-6 cells were transfected with the constructs using polyethylenimine (PEI) in Opti-MEM (Gibco) following standard protocols. After 72 hours, the supernatants were collected for analysis. Antibody concentration was quantified using an IgG2a ELISA kit (Bethyl Laboratories, E99-107). For antigen-binding assessments, rOspA (produced by GenScript) was coated onto plates at a concentration of 5 μg/mL. Binding was detected using the IgG2a Bethyl Laboratories ELISA kit (E99-107). Full-length LA-2 (produced by GenScript) was used as a positive control in both assays, diluted to 100 μg in 100 μL.

### Validation and Guide Testing for Full Length LA-2 Design

Four CRISPR-modified Hepa 1-6 cell lines were generated to validate the full-length LA-2 antibody design and assess the efficiency of different CRISPR guide RNAs. Each cell line integrated the full-length LA-2 IgG2a construct along with a puromycin resistance marker to enable selection of LA-2-positive populations. Hepa 1-6 cells were co-transfected with the LA-2 IgG2a construct and one of four px330 plasmids encoding SpCas9 and the following guide RNAs, targeting a site approximately 300 bp proximal of the albumin enhancer:

gRNA 1: CAGCTAACCTTCTGTCCTAG

gRNA 2: GCCTTAGCCAGTGTTTGCAC

gRNA 3: TCCTGTGCAAACACTGGCTA

gRNA 4: ACTGGCTAAGGCATGAACTT

Transfections were performed using Lipofectamine 2000 (Thermo Fisher) in Opti-MEM (Gibco), following standard protocols. Post-transfection, cells were selected with 2 µg/mL puromycin for three weeks to isolate stably integrated clones.

Total RNA was extracted from the CRISPR-modified Hepa 1-6 cells using the RNeasy Mini Kit (Qiagen), followed by cDNA synthesis with the Quantitect Reverse Transcription Kit (Qiagen). RT-qPCR was performed using the SensiFAST SYBR Hi-ROX Kit (Bioline), with mRNA levels normalized to β-actin and baseline expression in unmodified Hepa 1-6 cells. The following primer sets were used for amplification:

LA-2 mAb 1:

LA-2-1-qPCR-F: 5’-CTCCCTGTGGGTCTGAGTTT-3’

LA-2-1-qPCR-R: 5’-CCCATTGTTACATGCGTCGT-3’

LA-2 mAb 2:

LA-2-2-qPCR-F: 5’-TACCTGGTTGCAGGGTTGAT-3’

LA-2-2-qPCR-R: 5’-TCTGGCTTCATGCTCAATGC-3’

Albumin 1:

Alb-1-qPCR-F: 5’-CAAGAGTGAGATCGCCCATCG-3’

Alb-1-qPCR-R: 5’-TTACTTCCTGCACTAATTTGGCA-3’

Albumin 2:

Alb-2-qPCR-F: 5’-TGCTTTTTCCAGGGGTGTGTT-3’

Alb-2-qPCR-R: 5’-TTACTTCCTGCACTAATTTGGCA-3’

Puromycin resistance:

Puro-1-qPCR-F: 5’-CCACACCTTGCCGATGTC-3’

Puro-1-qPCR-R: 5’-CACCGAGCTGCAAGAACTC-3’

β-Actin:

Bact-1-qPCR-F: 5’-GGCTGTATTCCCCTCCATCG-3’

Bact-1-qPCR-R: 5’-CCAGTTGGTAACAATGCCATGT-3’

Supernatants were collected 72 hours post-transfection, and IgG2a levels were measured using the IgG2a ELISA kit (Bethyl Laboratories, E99-107) to assess antibody secretion.

### Generation of Full Length LA-2 expressing *Mus musculus*

Pronuclear injections were performed by the Whitehead Institute GEM Core. A vector containing the LA-2 expression cassette, without a selectable marker, was microinjected into BDF1 mouse embryos along with SpCas9 and guide RNA 3 (gRNA 3 sequence: TCCTGTGCAAACACTGGCTA), targeting a region 300 bp upstream of the albumin enhancer. Successfully injected embryos were implanted into pseudopregnant females. Of the 45 pups born, two carried the full-length LA-2 knock-in, and sequencing confirmed correct integration at the target locus in one of these mice.

### Genotyping Full Length LA-2 expressing *Mus musculus*

Genomic DNA was extracted from ear punches of LA-2-expressing *Mus musculus* using the GenElute Mammalian Genomic DNA Mini-prep Kit (Sigma-Aldrich) according to the manufacturer’s protocol. Genotyping was performed via PCR using PrimeSTAR® DNA Polymerase (Takara Bio) with two primer sets: one set amplifying the LA-2 heavy chain [LA-2-Heavy-F (5’-CCCATTGTTACATGCGTCGT-3’) and LA-2-Heavy-R (5’-AGGCATGAAGTCGGTTACCA-3’)], and the other set targeting the wild-type Albumin locus [Alb-F (5’-GCCTCTAATTCCCGTGTTCC-3’) and Alb-R (5’-TTGAACAGCCCACGAGAGAC-3’)]. PCR products were analyzed by agarose gel electrophoresis to assess zygosity. Homozygous mice, with complete integration of the LA-2 construct at both alleles, exhibited no amplification with the Albumin primers due to the size of the PCR product. Heterozygous mice displayed amplification from both sets of primers.

### *Mus musculus* husbandry

*Mus musculus* (C57BL/6) were maintained under 12:12 LD cycle of ∼400 lux (light) to <1 lux red light (darkness), with lights on from 6am to 6pm. Food and water were available ad libitum.

### Testing Mouse Serum for Full-Length LA-2

Serum samples were obtained from mice by collecting blood in BD Microtainer serum separation tubes. ELISA plates were coated with rOspA at 5 µg/mL. Serum samples were applied to the coated plates, and LA-2 antibody levels were detected using the IgG ELISA detection kit (Bethyl Laboratories, E99-131). Purified LA-2 IgG2a (produced by GenScript) was used as a standard.

### Bio-layer interferometry (BLI) assay

Kinetic analysis was performed using an Octet Red96 instrument following manufacturer’s instructions. Briefly, biotinylated OspA proteins were immobilized to streptavidin (SA) biosensors. The antigen-immobilized SA biosensors were then dipped into wells containing serially diluted (3.7-300 nM) antibody samples for 180 s for association. The sensors were then dipped into a kinetic buffer (PBST supplemented with 0.1% bovine serum albumin) for a 600 s dissociation step. Naked sensor was used as a non-specific binding control. Octet data analysis software (version 10.0.0.5) was used for kinetic curve fitting using the global fitting method.

### Linker Sequence Testing

Fifteen distinct linker sequences were evaluated to optimize expression, stability, and binding of the LA-2 scFv-Fc construct. Linkers were positioned between the heavy and light chains of the antibody. Three constructs utilized multimers of the GGGGS pentapeptide, while the remaining linkers were designed to adjust flexibility and length by varying the ratio of glycine and serine residues.

Lenti-X 293T cells were transfected with the linker constructs using polyethylenimine (PEI) according to standard transfection protocols. Seventy-two hours post-transfection, supernatants were harvested for analysis. Antibody concentration was quantified using a mouse IgG2b ELISA kit (Bethyl Laboratories, E99-109). Antigen binding was assessed by coating plates with 5 µg/mL of rOspA (produced by GenScript), followed by detection using the same IgG2b ELISA kit. Full-length LA-2 IgG2b antibody (produced by GenScript) was used as a standard.

### Generation of LA-2 scFv-Albumin expressing *Mus musculus*

B6 mouse embryos were microinjected with SpCas9, guide RNA (sequence: ACTCCAGTCTTTCTAGAAGA), and an LA-2 scFv-albumin expression cassette targeted to the Rosa26 locus. The expression cassette contained a CAG promoter driving the expression of LA-2 scFv fused to mouse albumin, with a C-terminal His-tag for detection and purification. Pronuclear injections were conducted at MIT’s DCM Transgenics Core, and successfully injected embryos were implanted into pseudopregnant females. Out of 18 pups, sequencing analysis confirmed correct integration at the Rosa26 locus in one pup.

### Genotyping LA-2 scFv-Albumin expressing *Mus musculus*

Genomic DNA was extracted from ear punches of LA-2 scFv-albumin expressing *Mus musculus* using the GenElute Mammalian Genomic DNA Mini-prep Kit (Sigma-Aldrich) according to the manufacturer’s instructions. Genotyping was performed by PCR using PrimeSTAR® DNA Polymerase (Takara Bio) with two primer sets. The first set targeted the wild-type Rosa26 locus, utilizing Rosa26-F (5’-CTCTGAGTTGTTATCAGTAAGGGAGCTG-3’) and Rosa26-R (5’-CCTCCCATTTTCCTTATTTGCCCCTATTA-3’).

The second set amplified the LA-2 scFv transgene using LA-2-scFv-F (5’-AAGGTCTCAAAAGAATGGGTTGGATCAAT-3’) and LA-2-scFv-R (5’-GGTTGAAGTGTGCTGGTGTAGTGTATAAG-3’).

PCR products were analyzed by agarose gel electrophoresis to assess zygosity. Heterozygous mice showed amplification from both the Rosa26 and LA-2 scFv primers, while homozygous mice displayed amplification only of the LA-2 scFv transgene.

### Testing mouse serum for LA-2 scFv-Albumin

Serum samples were collected from mice using BD Microtainer serum separation tubes. ELISA plates were coated with rOspA (produced by GenScript) at a concentration of 5 µg/mL. After applying the serum samples to the coated plates, LA-2 scFv-Albumin levels were detected using the Mouse Albumin ELISA Kit (Bethyl Laboratories, E99-134). As a standard, LA-2 scFv-Albumin-His (produced by GenScript) was used.

### Tick challenges

*Borrelia burgdorferi* infected nymphs used for challenging mice were prepared by feeding larval *Ixodes dammini* (Tufts colony) on mice (*Peromyscus leucopus* or *Mus musculus*) infected by lowpass strain N40 (wildtype), routinely maintained by tick-mouse-tick transfer, as described (Fikrig et al. 1992). At each challenge, all mice were infected from a single vial of infected nymphs with 70% infection rate. Mice were bled for serum prior to infestation. At infestation, mice were anesthetized with ketamine/xylazine and 10 nymphs applied to the ears and nape of the neck of each mouse, which was held with a wire restraining tube loosely wrapped with a paper towel. Mice were liberated into standard shoebox cages held within a larger cage containing an inch of water, and provided rodent chow and water ad libitum. Engorged nymphs were collected from the water at 4-6 days after infestation, and stored in standard tick vials at 21C and 95% RH for 14 days when they were analyzed for evidence of infection.

### Tick infection assay

Engorged nymphs were held 14 days as described (Fikrig et al 1992) and homogenized in 75uL of PBS in microfuge tubes. The tubes were briefly centrifuged to pellet gross debris, 7.5uL of the supernatant from each tick was applied to slides (Cel-Line 30-968, HTC), and allowed to dry before fixation in 100% acetone for 10 minutes. Slides were stained by indirect immunofluorescence using a rabbit polyclonal immune serum against *B. burgdorferi s*.*l*.; bound antibody was detected by staining with Alexa Fluor 488 conjugated anti-rabbit IgG. All slides were examined using epifluorescence at X400, and individual ticks scored for intact morphology as well as relative density. A subjective scale of 0-4 was used, with 0 representing absence of spirochetes and 4 representing typical intact cells and dense infection. The mean of the scores for all ticks recovered from each mouse represented the overall intensity of infection.

### Mouse infection assay

Mice challenged with infected ticks were held for 21 days and bled for serum, which was analyzed for evidence of *B. burgdorferi* specific IgG antibody using the C6 peptide assay as described (Bacon et al. 2003), except that an endpoint EIA was used instead of kinetic EIA. Briefly, biotinylated C6 peptide was bound to avidin coated wells of a flat bottomed microplate (Immulon 2), blocked with 3% fish gelatin in TBS (TBS-G), and 1:100 dilutions of mouse sera in TBS-G were incubated in duplicate for 1 hour at 37C. Bound antibody was detected using AP-goat anti-mouse IgG (gamma chain specific, Sigma), with a p-nitrophenyl phosphate substrate. A minimum of 6 negative control sera (uninfected B6 or C3H sera) optical densities were analyzed and used to calculate a cutoff value (mean + 3 standard deviations of the negative controls). Antibody to C6 is a sensitive indicator of *B. burgdorferi* infection.

### Xenodiagnosis

To definitively determine mouse infection status after challenge, xenodiagnosis (Telford et al 2014) was performed. At day 21 after infected tick challenge, mice were infested with larval I. dammini ticks (Tufts colony), which were allowed to engorge. Engorged larvae were collected, placed into standard tick vials, and held at 21C and 95% RH until they molted 4-5 weeks later. After molting and hardening, samples from each vial were dissected and guts smeared onto slides. The slides were dried, fixed in acetone, and stained by indirect immunofluorescence using an immune rabbit polyclonal serum against *B. burgdorferi s*.*l*. with secondary antibody comprising AlexaFluor 488 goat anti-rabbit IgG. Slides were examined for a minimum of 50 fields (X320) before declaring a negative. Any tick containing spirochetes (e.g., even 1 of 5 ticks) is evidence that the mouse is infected and is infectious.

**Supplemental figure 1.**
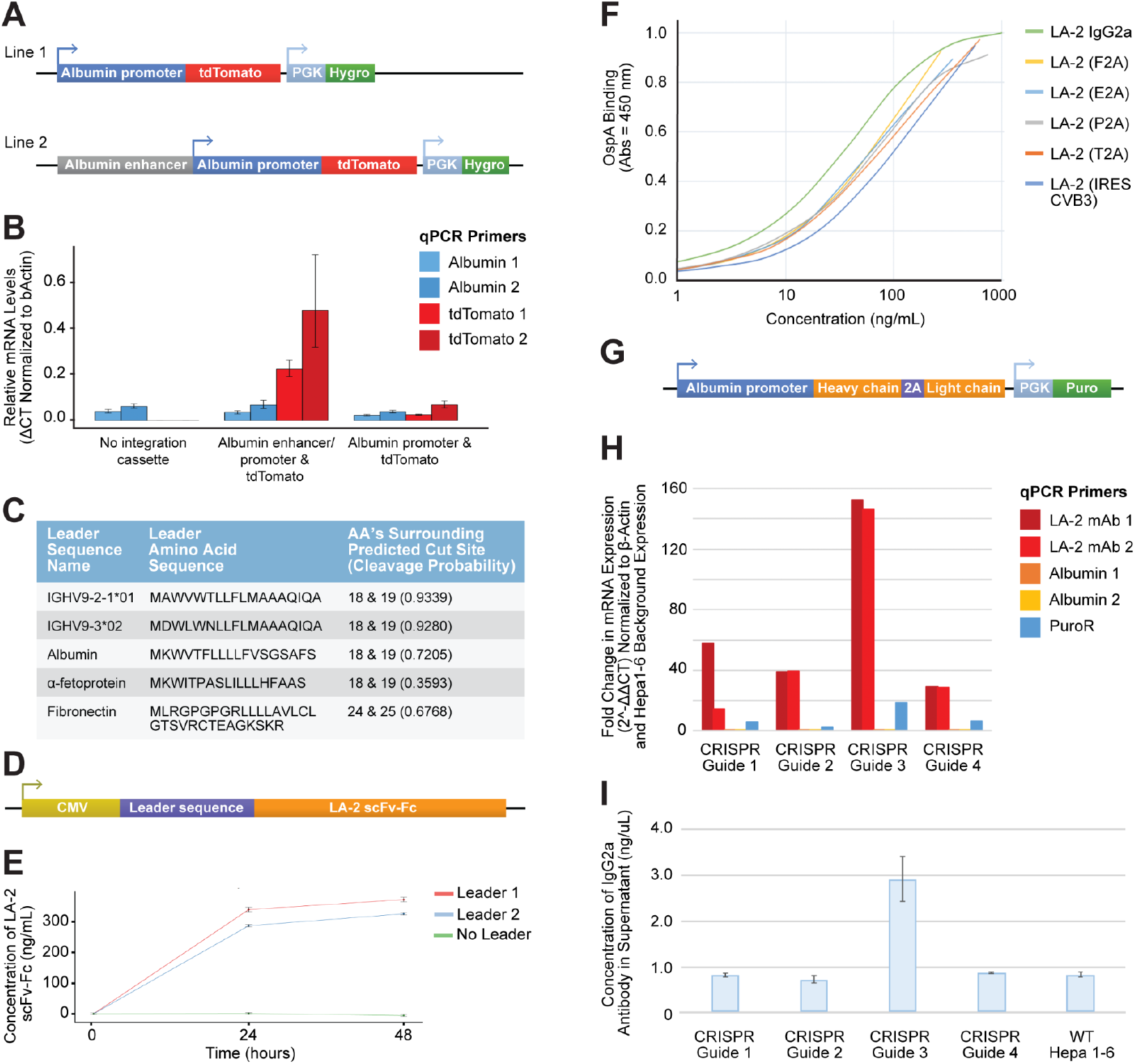
Optimization and Validation of LA-2 Antibody Expression in Hepa 1-6 Cells. **A)** Schematic representation of the CRISPR-modified Hepa 1-6 cell lines used for testing albumin expression machinery, incorporating the minimal albumin promoter with and without the albumin enhancer. **B)** RT-qPCR analysis showing relative mRNA levels of tdTomato and albumin in Hepa 1-6 cells transfected with the constructs described in (A). **C)** Leader sequences evaluated for their ability to direct LA-2 antibody secretion, including sequences from mouse albumin, alpha-fetoprotein, and fibronectin, along with their predicted cleavage efficiency. **D)** Schematic of the LA-2 scFv-Fc expression construct under the control of a CMV promoter, including the leader sequences tested for secretion efficiency. **E)** ELISA results comparing LA-2 scFv-Fc concentrations in the supernatant of Hepa 1-6 cells transfected with constructs containing two different leader sequences, measured over a 48-hour period. **F)** OspA binding activity of LA-2 antibodies produced using different bicistronic elements, measured by ELISA at various concentrations. **G)** Schematic of CRISPR-modified Hepa 1-6 cell lines expressing the full-length LA-2 antibody. **H)** RT-qPCR analysis showing fold change in LA-2 mRNA in engineered Hepa 1-6 cells with different CRISPR guides, normalized to β-actin and background Hepa 1-6 expression. **I)** Concentration of IgG2a in the supernatant of CRISPR-modified Hepa 1-6 cells, quantified by ELISA, demonstrating successful antibody secretion in cells engineered with guide 3.

**Supplemental figure 2.**
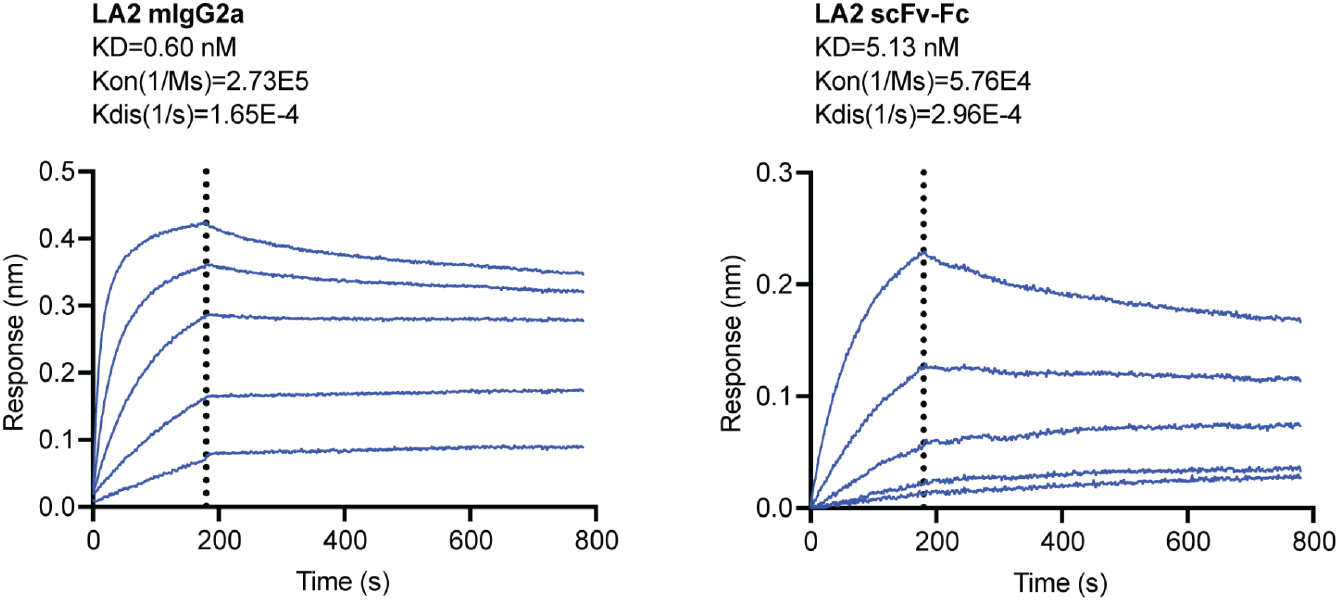
Binding Affinity of LA-2 IgG2a and LA-2 scFv. Binding affinity data for LA-2 IgG2a and its scFv counterpart with a Vh-(G4S)3-Vl linker. Association rate constant (kon), dissociation rate constant (koff), and equilibrium dissociation constant (Kd) values are reported.

**Supplemental figure 3.**
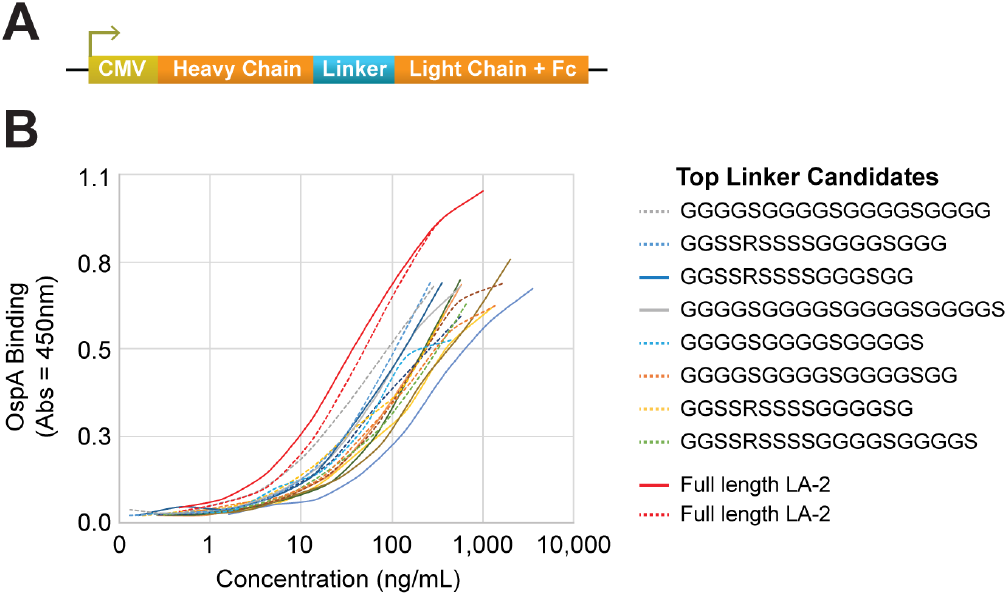
Evaluation of Linker Sequences for Optimizing LA-2 scFv-Fc. **A)** Schematic representation of the expression construct used to evaluate different linker sequences between the LA-2 scFv heavy and light chains. The construct includes a CMV promoter driving the expression of the LA-2 scFv-Fc fusion protein, with various linkers designed to modulate flexibility and length. **B)** OspA binding data from ELISA assays comparing the performance of different linker sequences in Lenti-X 293T cells 72 hours post-transfection. The binding profiles of the top linker candidates were assessed against purified full-length LA-2 antibody (red lines).

